# Character displacement in shorebirds

**DOI:** 10.1101/2025.09.02.673710

**Authors:** Lucyle Hernandez-Possémé, Glenn Le Floch, Vincent Bels, Nicolas Schtickzelle, Michel Baguette

## Abstract

Modification of phenotypic traits induced by character displacements allow species to coexist by reducing interspecific competition. This study explores if and how shorebirds partition feeding ressources by foraging at different water depths outside the breeding season. During this period, communities of up to dozen shorebird species coexist on suitable habitats at land-water interfaces. By video recording 22 species of migrating and wintering shorebirds in different sites, we show that their distribution according to the water height at which they forage is not random but follows a gradient of increasing water height according to increasing species body size. Species-specific beak and tarsus lengths are positively correlated with the water height at which the different species forage. Our data do not support a trade-off in investment between beak length and tarsus length in our study species set. Congeneric species that are expected to be morphologically and ecologically more similar forage at contrasted water heights. Altogether, our findings supports the idea of resource partitioning, and consequently of competition relaxation, by morphological and/or by behavioral character displacements.

## Introduction

How closely related species share resources, and thus potentially coexist, is one of the many questions in ecology (*e*.*g*., De León et al. 2014; Barnagaud et al. 2021; Seidelmann and Mostaghim 2025). When several species exploit similar resources (*i*.*e*., have overlapping ecological niches), whatever their nature, interspecific competition translates into selection pressure leading either to the competitive exclusion of one of the species (Gause 1934; Hardin 1960), or to the emergence of mechanisms allowing them to coexist (*e*.*g*., Losos 2000; Pfennig and Pfennig 2009). In this case, phenotypic trait modifications are implemented to promote niche partitioning and reduce interspecific competition. Character displacement describes this process, whereby morphological, behavioral or physiological traits diverge between sympatric species to minimize niche overlap (Brown and Wilson 1956; Schluter 2000). This process is well documented in several taxa, particularly birds, with the emblematic example of Galapagos finches (Lack 1947; Grant and Grant 2024). In these birds, repeated measurements from year to year of the beak size and shape of two sympatric granivorous species show that these parameters evolve rapidly with the size of the seeds available, in order to minimize niche overlap (Grant and Grant 1993, 2006, 2024). There is therefore a close relationship between the resource (seed size), the morphological trait used for its exploitation (beak size and shape) and the fitness of individuals (survival and reproduction of those best adapted to the available seeds).

Migratory birds, particularly shorebirds (Scolopaci and Charadrii), are also a recognized example of food resource partition. Outside the breeding season, these birds feed mainly at land-water interfaces in fresh, brackish, or marine waters. During their migratory stopovers or on their wintering grounds, dozen different species are found in sympatry, sharing space and the available food, mostly consisting of invertebrates and plant material (*e*.*g*., Cramp and Simmons 1983; Colwell 2010). Despite this co-occurrence, direct competition appears to be limited (*e*.*g*., Bocher et al. 2014; Novcic 2016). Variation in the size of prey consumed, foraging schedules and preferred microhabitats would facilitate coexistence (Bocher et al. 2014; Bellefontaine and Hamilton 2023). The contrast found in morphological structures related to prey capture (*e*.*g*., beak, legs) (*e*.*g*., Barbosa and Moreno 1999; Durell 2000; van der Kam et al. 2004; Bellefontaine and Hamilton 2023) could therefore be a response to competition between these different species. Differences in beak length and shape and differences in leg length would influence the trophic niches of shorebirds by adapting the nature of the prey caught and the height of water and sediment at which they feed (Zwarts and Blomert 1992; Barbosa and Moreno 1999; van der Kam et al. 2004; Nebel et al. 2005; Bocher et al. 2014). While this explanation seems satisfactory, as it corresponds to the theoretical principle of character displacement defined above, it lacks quantitative validation to date.

The aim of our work is to add to this knowledge by documenting any character displacement in migrating and wintering shorebirds. We worked on some twenty migrating and wintering Western Palearctic shorebird species, in different localities to avoid topological constraints on our study. We first determine the height of water at which they feed. Under the hypothesis of character displacement, we expect (1) the species to be distributed along a gradient of water height, and (2) the existence of a positive relationship between their beak and leg lengths, on the one hand, and the height of water at which they forage on the other. Finally, as beak and leg lengths may have followed differential evolutionary trajectories among species, the existence of a possible trade-off in investment between these two morphological traits will also be investigated.

## Materials & Methods

### Data collection

The foraging and feeding behaviors of shorebirds were video recorded opportunistically at several sites in France and Belgium (Western Europe) between 2022 and 2025 (S1). Observers filmed while positioned at a distance between 3 and 50 m of focal individuals to avoid disturbing the birds.

The video sequences were recorded using an OMD-EM1x camera equipped with a M.Zuiko Digital Pro ED 150-400 mm F4.5 lens, providing up to 40 X magnification. Focal individuals (Altmann, 1974) were selected (in groups when necessary) and filmed at a speed of 25 or 60 frames per second. Data collection was carried out on 22 species of shorebirds observed in the study area, for which we have at least 5 individuals per species, divided into two of the three suborders of Charadriiiformes: 15 species of Scolopaci and 7 of Charadrii (Černý & Natale, 2022).

### Morphological and behavioral data

Locomotion behaviors (how individuals move around their feeding habitats) and capture behaviors (how individuals catch and locate their prey) were categorized based on a recent repertoire (Baguette et al., 2024). There are five capture behaviors (1: pecking, 2: probing, 3: pecking-probing, 4: sweeping, 5: routing) and three locomotion behaviors (1: run-stop-run, 2: continuous walking, 3: continuous walking-continuous swimming).

In addition, aggregated data at the species level were used for several morphological variables (longest toe length, wing length, tarsus length, beak length, weight, and size) and for each species studied. These data were extracted from the synthesis by Cramp & Simmons (1983). Each value represents a species average, *i*.*e*., an estimate of the average trait for all individuals of a species, regardless of sex.

### Quantifying foraging water height from video recordings

We analyzed the videos using Kinovea software (https://www.kinovea.org, v. 2023.1) to quantify the exposed part of the legs (*i*.*e*, the length of the legs out of the water for each measurement), for each focal individual selected on a video at regular intervals (every 15 seconds). The species tarsus and beak length values (mm) were measured simultaneously to serve as a scaling value and convert the exposed leg length from a relative (in pixels) to an absolute measure using their known average value for the species. In addition, the tibia length was measured in each video. We computed the length of the leg submerged in the water by subtracting the exposed leg length from the total length of the leg (tarsus plus tibia length). The data were then averaged to obtain a water depth value per individual.

### Statistical analyses

First, we constructed a boxplot (from R package GGPLOT2) of the water depth used by the 22 species. Given the strong correlation between the variables of interest (tarsus and beak length : R= 0.66, n=22, p < 0.001), we constructed three body size indices for each species based on the various morphological variables mentioned above, on which we performed phylogeny constrained PCAs (Principal Component Analyses, *phyl*.*pca* function from R package PHYTOOLS) to control for the effect of evolutionary relatedness among species (the phylogenetic tree used for this method comes from Černý & Natale, 2022, with the 22 target species). Three PCA were performed: one without tarsus length (tarsus body size index, BSIT), another without beak length (beak body size index, BSIB), and the last with all morphological variables (total body size index, BSI_total). In each case, the first principal component (PC1) was extracted from the phylomorphospace, with the coordinate of each of the 22 species on this axis defining the size indices. The BSI_total index was used to analyze the water depth at which the species feed. The next step was to perform a linear regression (*lm* function from R package STATS) for BTSI and BBSI to assess the relationship between PC1 and the excluded variable (the tarsus or beak, respectively). The residuals from these regressions were then extracted and interpreted as indices of morphological deviation specific to each species, relative to the values expected given their overall morphology (RTE: Relative To Expected).

Linear regressions were performed to explore the relationships between these residuals and the water depth that the species should use based on its body size (RTE_water). Finally, a trade-off in the investment in beak lengths and leg lengths within our study species was sought by performing a linear regression between the RTE of the tarsus and the RTE of the beak.

## Results

### Species distribution as a function of water height

The distribution of the species studied according to the water height at which they forage is not random but follows a gradient of increasing water height (Fig. 1). Species belonging to the two suborders Scolopaci and Charadrii are not arranged in the same way along this gradient: the distribution of Scolopaci is monotonic, while that of Charadrii is bimodal (Fig. 1A, 1B). Among the Charadrii, one group is made up of Plovers (Cha_hia, Thi_dub, Och_ale, Plu_squ) and Oystercatcher (Hae_ost), which feed at low water levels, and another group with Pied avocet (Rec_avo) and Black-winged stilt (Him_him), which frequent much higher water levels (Fig. 1A, 1B). The classification of locomotion and capture behavior for each species according to water level is not random either (Fig. 1A, 1B). For locomotion, run-stop-run behavior is only present in species that feed at low water levels, while locomotion by continuous walking and swimming is only found in one species with one of the highest water levels. Continuous walking, present in the majority of species, is used at all water heights. Capture behavior also shows a certain organization according to water level. Pecking and routing are found in species that feed at low water levels, in contrast to sweeping at high water levels. Combined pecking and probing behavior is relatively homogeneous, used at several different water depths, while exclusive probing tends to be observed at medium to high water depths. A comparison of the water depths frequented by *Calidris* species (Fig. 1C) shows that two species are found at the extremes of the distribution, with the Purple Sandpiper (Cal_mar) at zero water depth and the Curlew sandpiper (Cal_fer) at high water depths. The other four species are present at low to medium water depths, with a gradient from the Sanderling to the Ruff (Cal_pug). A similar pattern is observed for the genus *Tringa* including the Common Sandpiper (Act_hyp) (Fig. 1D).

**Figure 1.**
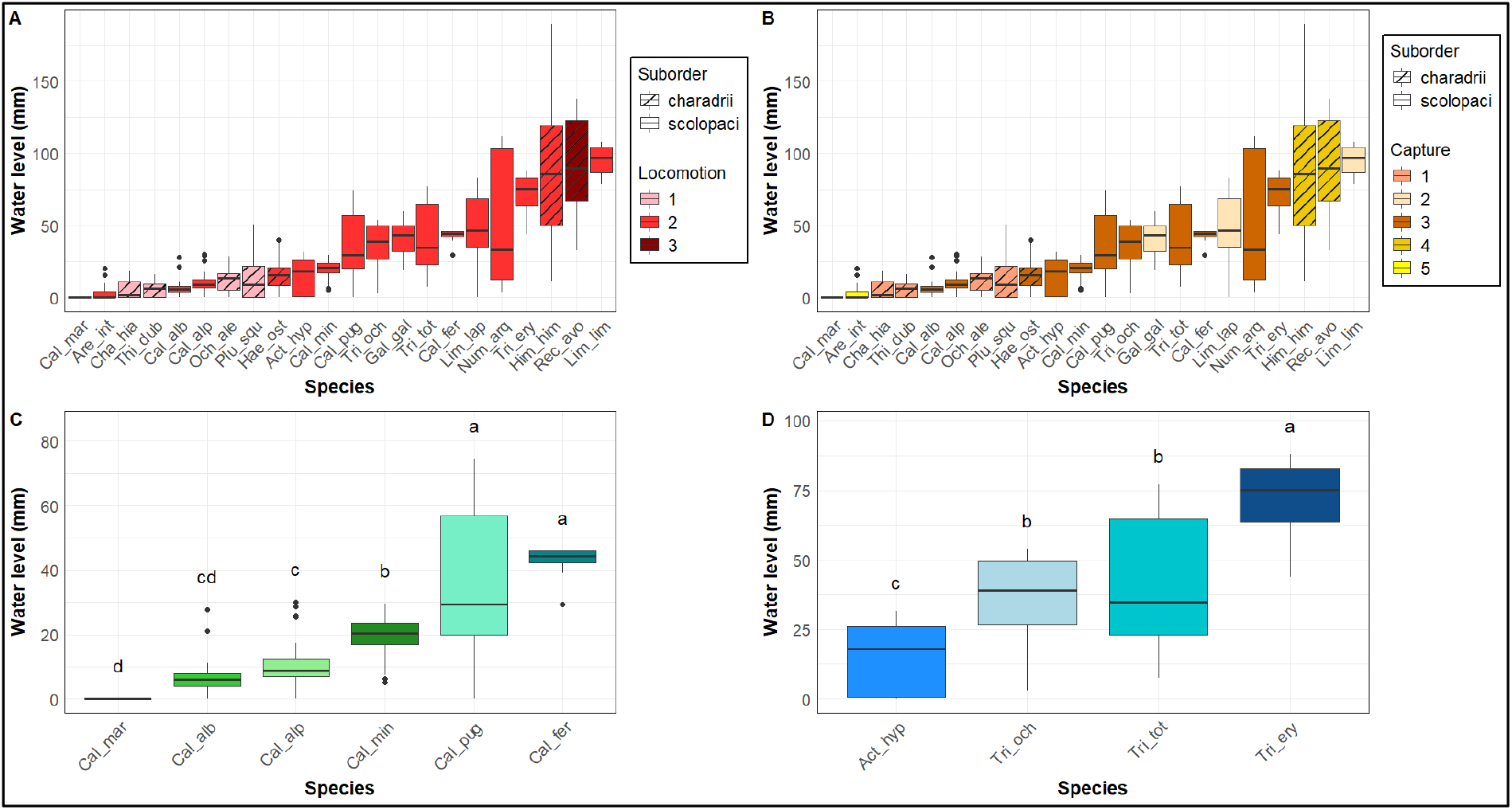
Distribution of species according to the water level at which they feed. (A&B) According to suborders and respectively locomotion (1: run-stop-run, 2: continuous walking, 3: continuous walking-continuous swimming) and capture behaviors (1: pecking, 2: probing, 3: pecking-probing, 4: sweeping, 5: routing). (C&D) Distribution of species belonging of the genus, *Calidris* and *Tringa* respectively. Different letters indicate significant differences between groups (Tukey HSD test, p < 0.05).

### Relationship between species morphological characteristics and water level

The height of water used by our study species can be predicted from the body size index as computed from morphological variables (Fig. 2): the height of water used by a species is a linear, positive function of its body size. Given that larger species feed in deeper water and larger species tend to have longer legs and longer beaks, the interesting question becomes: what if we control for body size, for both the water depth and the leg or beak length ?

**Figure 2.**
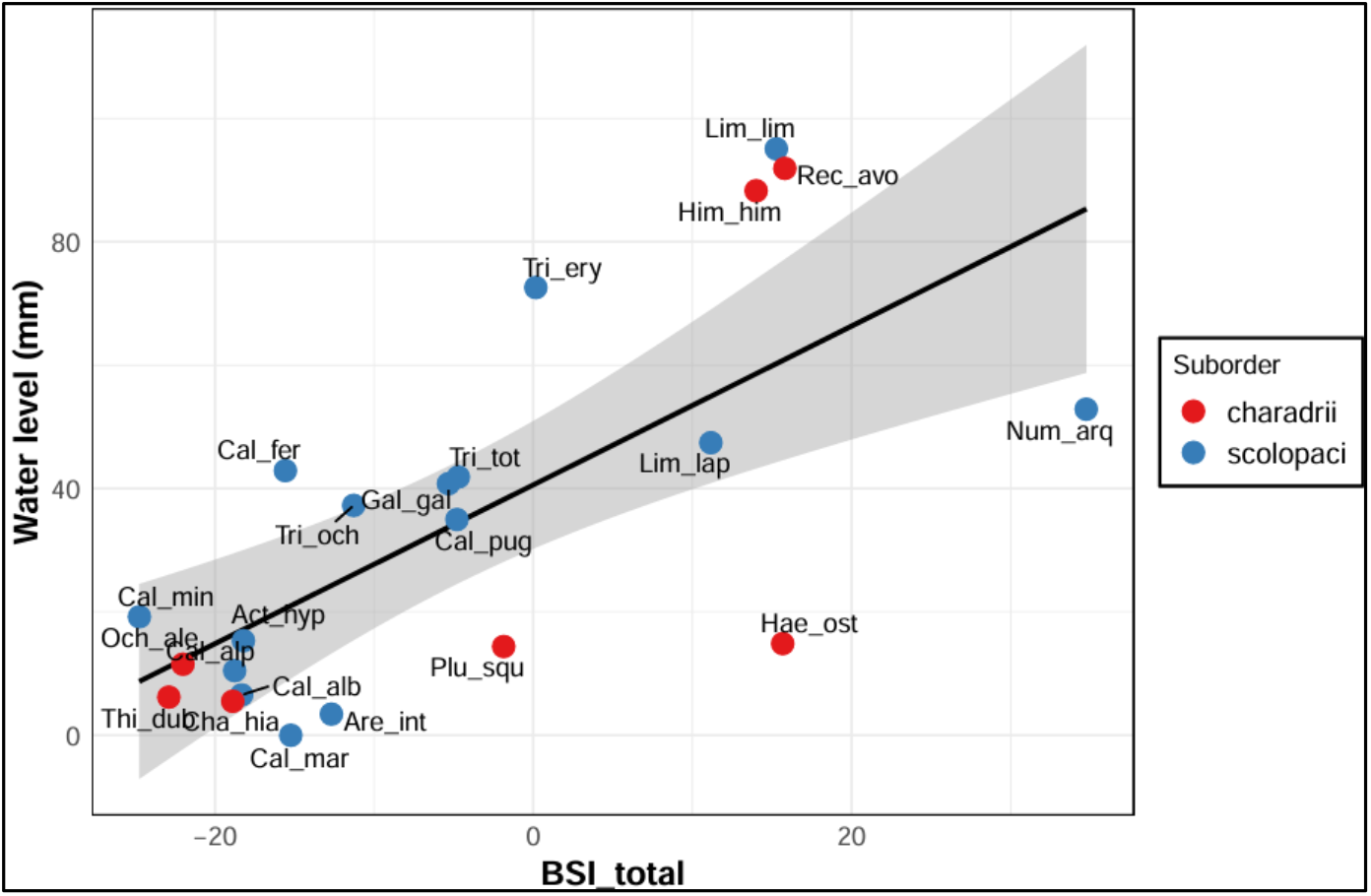
Relationship between water depth and BSI_total, the Body Size Index for all morphological variables (R^2^ = 0.5, p < 0.001***).

The RTE for the water level is significantly and positively related to the RTE of tarsus and beak lengths (Fig. 3A, 3B).

**Figure 3.**
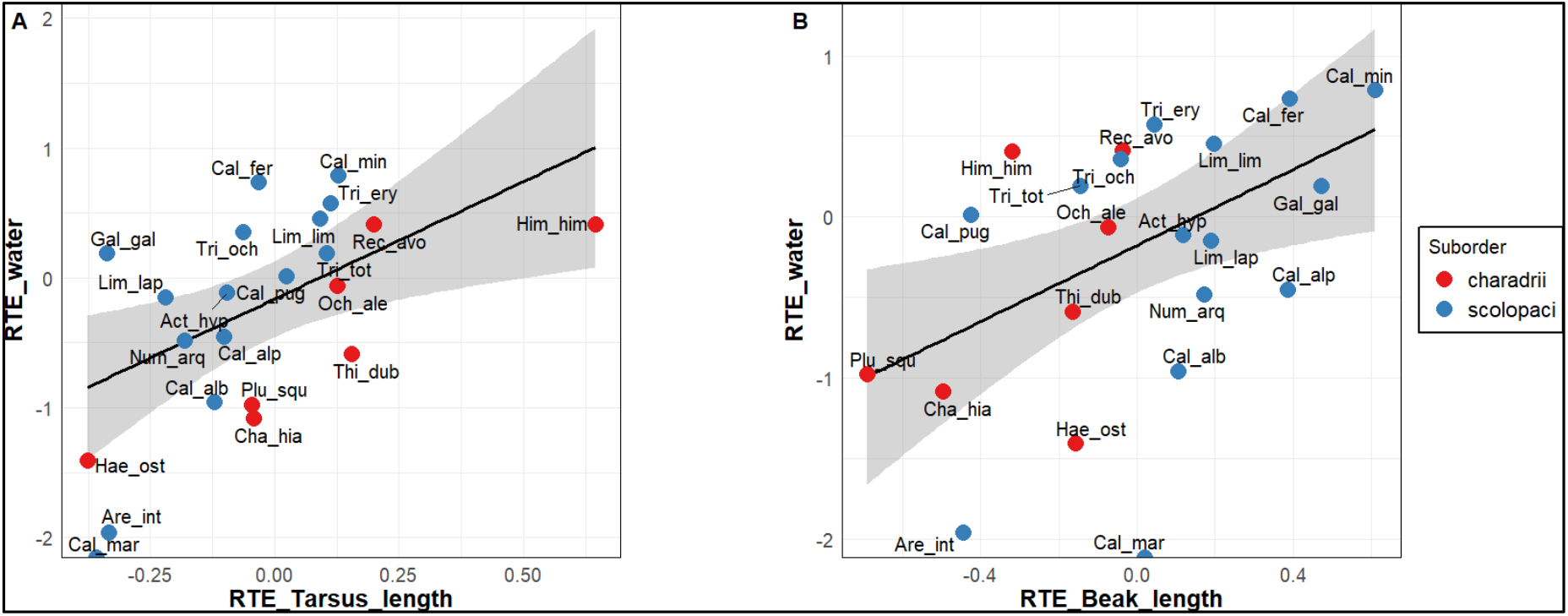
Relationship between the deviation of actual values of water height used by each species from its predicted value according to their body size index (RTE_water), and the deviation of their tarsus (RTE_Tarsus_length (A)), and beak lengths (RTE_Beak_length (B)) actual values from their predicted value according to their body size index. ((A) R^2^ = 0.37, p = 0.003 **). ((B) R^2^ = 0.19, p = 0.04*).

### Relative investment in tarsus length and beak length

We expected a negative relationship under the hypothesis of a trade-off at the species group level. We found a very slight indication of it (Fig. 4) but the relation is not statistically significant because of the large variation among species, which is a good illustration that different strategies exist: some have longer RTE tarsus length, some longer RTE beak length. Some species have a longer beak than expected, such as the Common snipe (Gal_gal) with a much smaller tarsus. Other species have a much shorter beak and tarsus than expected, such as the Ruddy turnstone (Are_int). Still others, like the Black-winged stilt (Him_him), have large tarsi but small beaks. No species has two strongly positive values.

**Figure 4.**
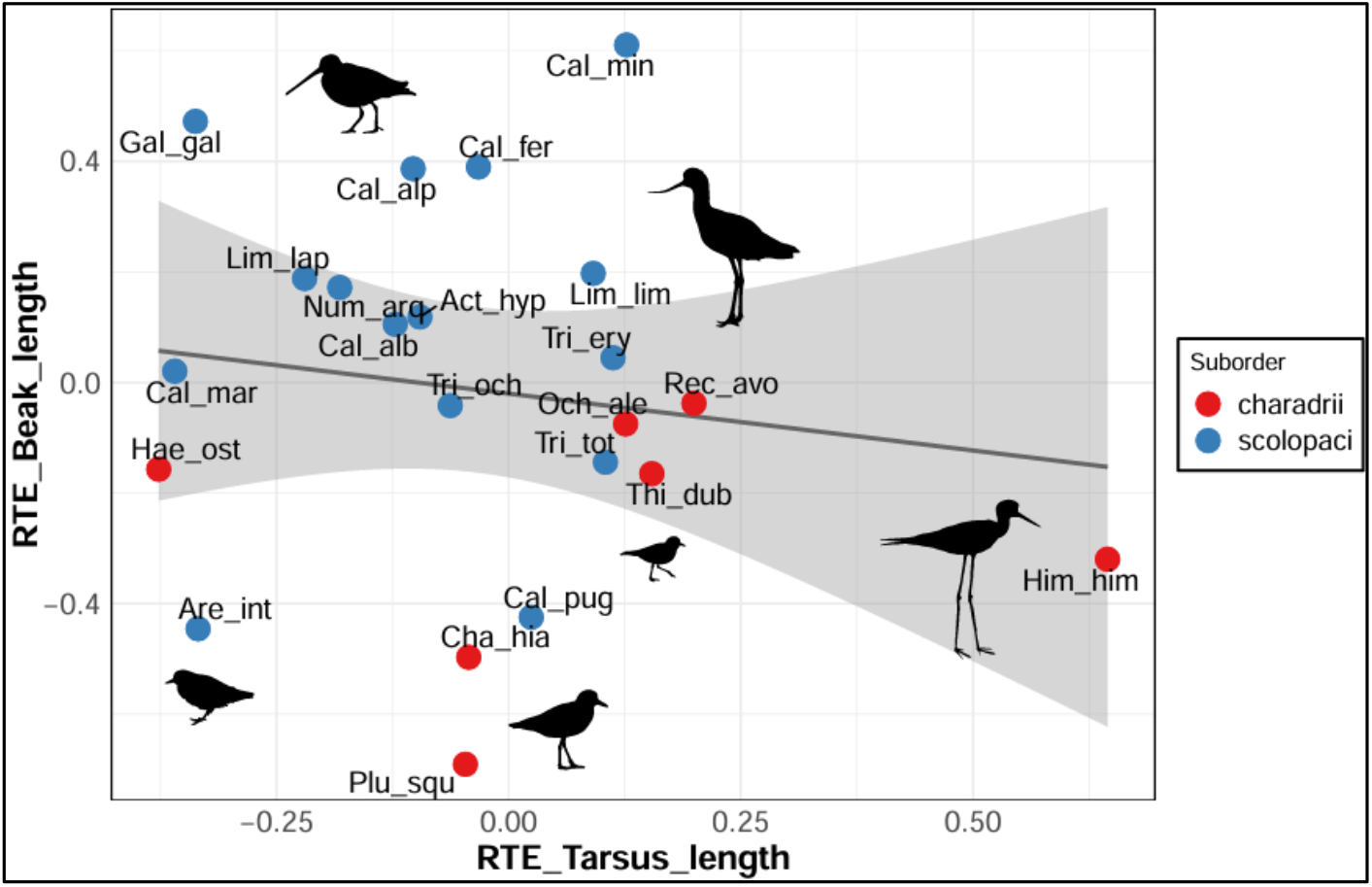
Relationship between the deviation of the beak (RTE_Beak_length) and tarsus length (RTE_Tarsus_length) actual values from their respective predicted value according to their body size (R^2^ = 0.02, p = 0.52).

## Discussion

Our results highlight that the spatial distribution of shorebirds based on the water depth at which they forage is not random. Some species or taxonomic groups (suborders) tend to aggregate preferentially at specific water depths. This structure suggests an ecological partitioning between species (*e*.*g*., Colwell and Landrum 1993; Catry et al. 2016) and confirms previous works. Baker and Baker (1973) indeed studied the water depth at which six species of shorebirds forage and used this parameter to define microhabitats. Barbosa and Moreno (1999) also looked at water depth during foraging and noted a difference between the two suborders of shorebirds, with Scolopaci feeding at deeper water depths than Charadrii, with the exception of Recurvirostridae (Pied Avocet and the Black-winged Stilt). Recently, the work of Yu et al. (2023) in China analyzes the use of foraging microhabitats in four species of shorebirds and shows that water depth is positively related to beak and leg length. A similar analysis was conducted by Aarif et al. (2024) in India and indicates that water depth is positively related to beak length, but not leg length. However, these two studies are limited by the small number of species studied in the first case and by methodological problems in the second, as water depth is not measured directly, which probably explains why tarsus length is not correlated with water depth, contrary to what we observe here and according to other studies (*e*.*g*., Barbosa and Moreno 1999; Yu et al. 2023).

We observe differences between suborders: Charadrii feed in shallow water (except for the Pied Avocet and the Black-winged Stilt), while Scolopaci have no preferred water depth and frequent all water depths. Besides, Charadrii most often frequent shallower water depths than expected based on the length of their tarsus and their beaks and vice versa for Scolopaci. According to the dated phylogeny of Černý and Natale (2022) our sample of 7 species of Charadrii is composed of 4 species with short beaks belonging to Charadriiidae (plovers), an ancient clade of the sub-order, and 3 species with longer beaks belonging to a lately diverged clade. The variations observed between the suborders could be explained by their foraging techniques: Scolopaci are able to locate food by touch, unlike plovers, which rely on sight (*e*.*g*., Barbosa and Moreno 1999; Thomas et al. 2004). These techniques can be linked to different modes of locomotion and capture behavior (Baguette et al. 2024), which, as we show here, are practiced at contrasting water depths.

A remarkable result is the non-significant relationship between the deviation from the expected value of the beak and that of the tarsus based on body size, suggesting the absence of an evolutionary trade-off whereby species tend to invest in either a long beak or a long tarsus, but not in both. The resulting wide morphological variation could explain why these species have extremely diverse diets, ranging from invertebrates and plant material buried deep in the sediment to preys captured at the surface. In areas where several species feed at similar water depths, these morphological differences are expected to reduce interspecific competition (*e*.*g*., Stillman et al. 2000; Piersma 2007).

The morphological differences we highlight here could be the result of character displacement, a process recognized as an important driver of ecological differentiation and niche diversification. Although some species of shorebirds feed at the same water depths, their morphology differs (beak and/or tarsus length). Moreover, congeneric species that have a common evolutionary trajectory are expected to be morphologically and ecologically more similar (Darwin 1859) and thus have a great niche overlap. We show here that within the genus *Calidris* and *Tringa*, species forage at contrasted water depth, which supports the idea of resource partitioning, and consequently of competition relaxation. This partitioning has already been documented in *Calidris*, where phylogenetically close species exploit different microhabitats for foraging (Zwarts et al. 1990; Nebel et al. 2005) which thus translates either in morphological and/or or behavioral character displacements. Such differentiation would allow coexistence without direct competition, with specialization on different prey and/or specific foraging modes, as proposed by Tilman (1977). The endpoint of this process is what we observe here, i.e., either the sharing of a common foraging niche by targeting different resources (Schluter 2000) or the use of the same resource within different microhabitats (Zwarts et al. 1990; Nebel et al. 2005). These two potential issues to the same problem of resource partitioning demonstrate how it is important to place each species in its ecological and evolutionary context (Losos 2000).

In conclusion, our work has shown that the distribution of shorebirds according to water depth is not random and is linked to their morphology. We suggest that variability in tarsus and beak lengths allows for species specialization, thereby limiting competition between them. The absence of an evolutionary trade-off between traits highlights the diversity of feeding strategies used to coexist in the same habitat. It should be noted here that intra-specific variability, *i*.*e*., within the same species, is not taken into account. We know that this variability exists with significant sexual dimorphism in the species studied (Nebel and Thompson 2011). This variation would limit competition for resources between males and females. However, as the objective of this study is to compare species-specific relationships between habitat use and morphological traits, we focused here on species and not on individual. Intra-specific variability was therefore not considered.

The consideration of this intra-specific variation in morphology and its role in microhabitat selection by different species could, however, constitute an interesting research avenue for improving our study. Besides, the inclusion of external variables such as the nature of the substrate on which shorebirds forage could also be interesting to consider in order to observe if it has an influence on microhabitat selection. Despite these limitations, our study provides new information on the links between food acquisition behavior and morphology, and more broadly on the role of character displacement in promoting species coexistence.

## Supplementary materials

**S1.**
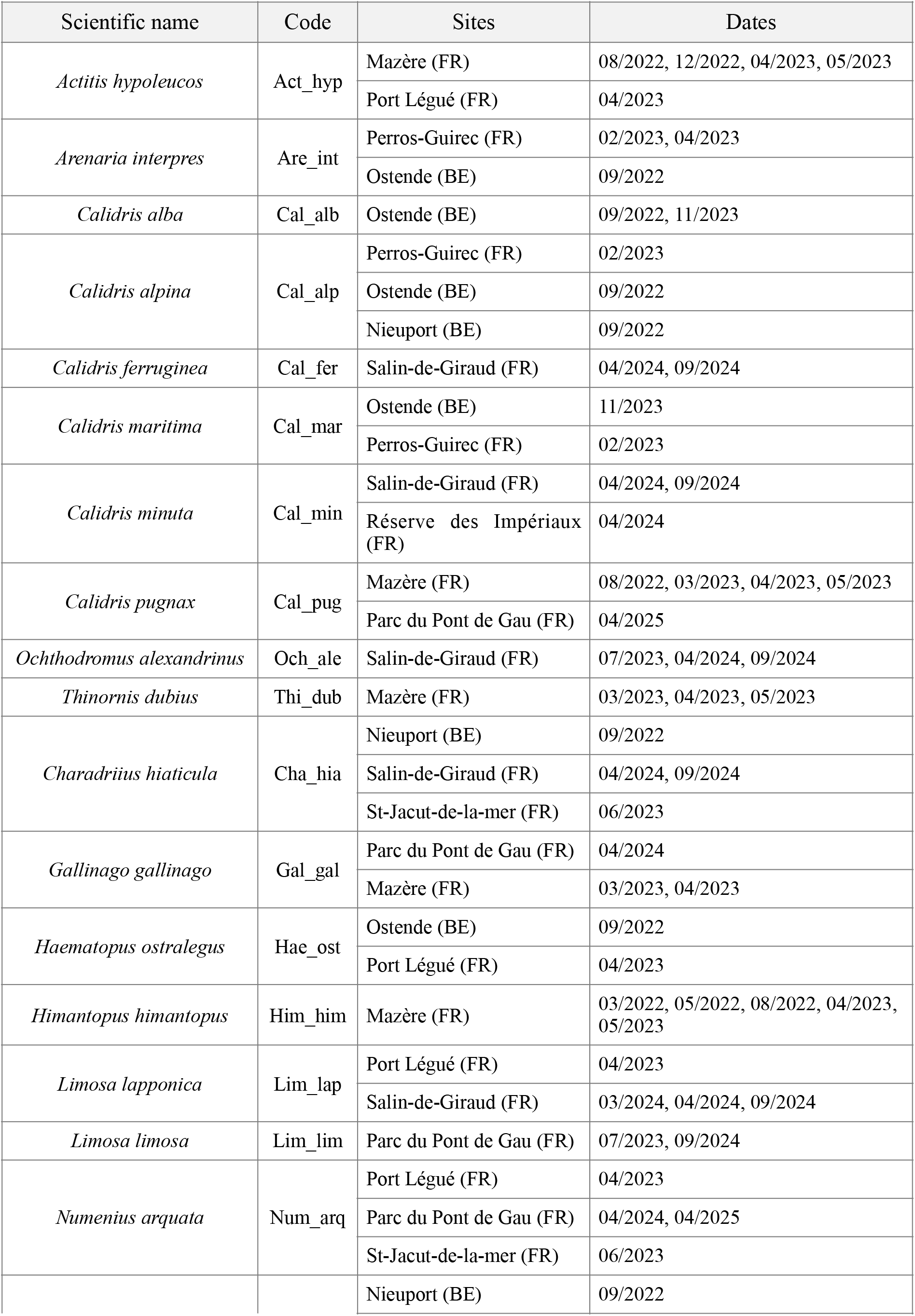
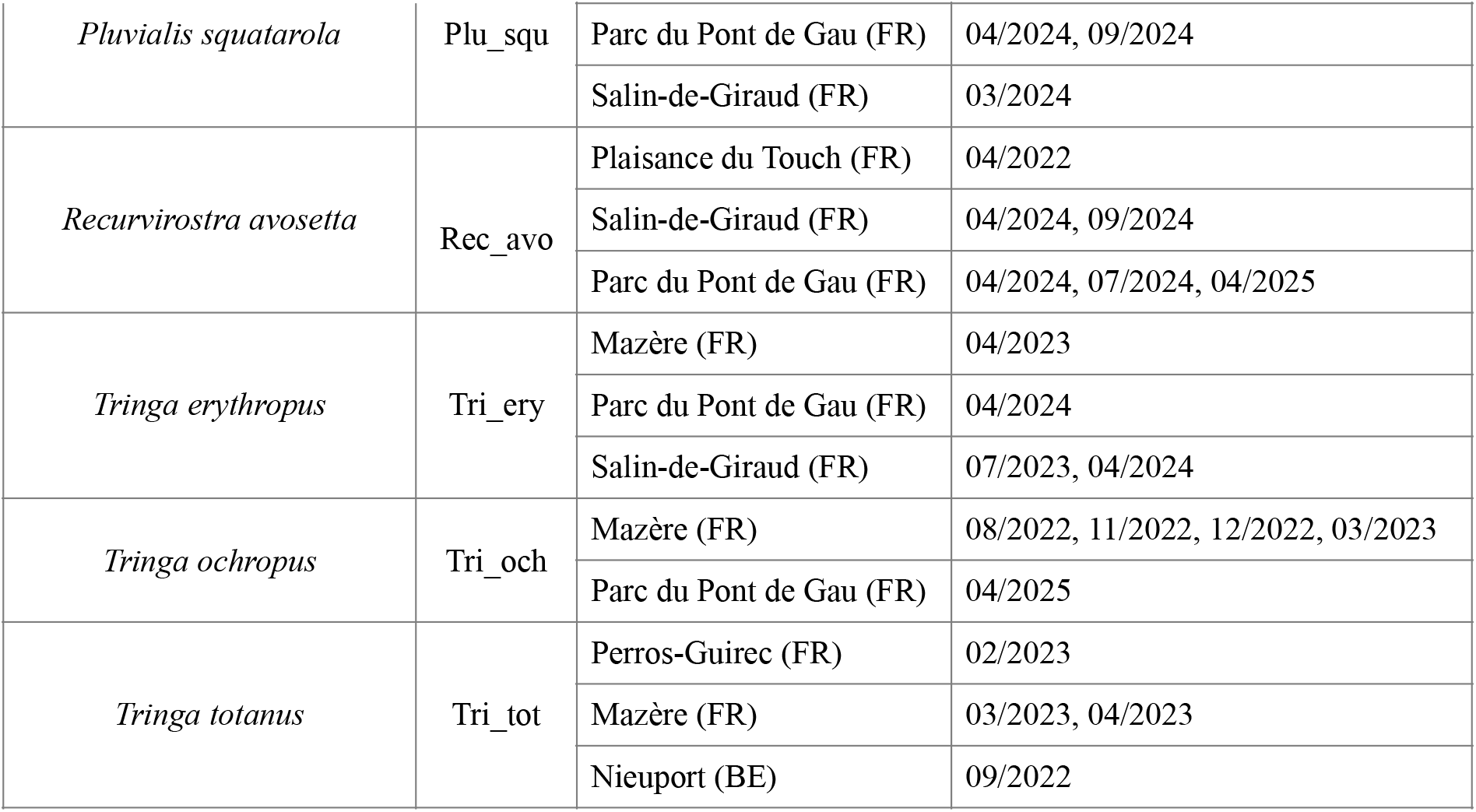
Sites, sampling dates and codes used for every species in the analyses.

